# TDP-43 subcellular mislocalisation is correlated with loss of optineurin binding for frontotemporal dementia and amyotrophic lateral sclerosis associated *TBK1* missense variants

**DOI:** 10.1101/2022.06.02.494603

**Authors:** Lisa J. Oyston, Lauren M. Boccanfuso, Lauren Fitzpatrick, Johnny Zhang, Marianne Hallupp, John B. Kwok, Carol Dobson-Stone

## Abstract

**Background:** Frontotemporal dementia (FTD) is one of the most common forms of younger-onset dementia. FTD is genetically, pathologically and clinically related to amyotrophic lateral sclerosis (ALS), a rapidly progressive neurodegenerative disorder. Mutations in TANK-binding kinase 1 (*TBK1*) have been identified as a rare cause of FTD and ALS. TBK1 has known roles in inflammation and autophagy and interacts with other FTD and ALS proteins such as optineurin (OPTN): however, which of its roles are important to FTD/ALS pathogenesis remains undetermined. To date, >90 *TBK1* rare variants have been identified in FTD/ALS patients: >50% of these are missense variants of unknown significance (VUS).

**Methods:** In this study, we have used a functional assay pipeline to investigate the effect of 16 *TBK1* VUS with *in-silico* evidence of pathogenicity, together with two known pathogenic mutations and a common benign *TBK1* polymorphism. Our assay pipeline evaluated the effect of *TBK1* VUS on steady-state levels of TBK1, kinase activity and binding to OPTN. We also assessed the impact of *TBK1* VUS on a key neuropathological feature of FTD and ALS cases: mislocalisation of neuronal TDP-43 from the nucleus to the cytoplasm.

**Results:** We observed some *TBK1* VUS that had similar effects to *TBK1* loss-of-function mutations, demonstrating decreased kinase activity and loss of OPTN binding. Both known pathogenic mutations and several *TBK1* VUS also increased the cytoplasmic/nuclear ratio of TDP-43 and this inversely correlated with their degree of OPTN binding but not with kinase activity.

**Conclusions:** These results suggest that loss of the direct interaction between TBK1 and OPTN is more critical to FTD and ALS pathogenesis than TBK1’s kinase activity. However, further studies are needed to elucidate exactly how loss of TBK1 binding to OPTN leads to TDP-43 pathology and ultimately neurodegeneration.

## Background

Frontotemporal dementia (FTD) is one of the most common forms of younger-onset dementia and involves the progressive degeneration of the frontal and temporal lobes of the brain. Increasing genetic evidence and clinical observations have linked FTD to amyotrophic lateral sclerosis (ALS) [1]. ALS is a progressive neurodegenerative disorder that affects the upper and lower motor neurons leading to muscle weakness and paralysis. 40% of FTD cases and 10% of ALS cases are familial, suggesting a strong genetic component to these diseases. Several genes have been identified where mutations can lead to either or both FTD and ALS (FTD-ALS genes), with the most common cause being a GGGGCC hexanucleotide repeat expansion in *C9orf72*, accounting for 11-13% of cases [2]. Sequestosome 1 (*SQSTM1*), valosin containing protein (*VCP*), coiled-coil-helix-coiled-coil-helix domain containing 10 (*CHCHD10*), optineurin (*OPTN*) and the recently identified TANK-binding kinase 1 (*TBK1)* are other significant genetic contributors to the FTD-ALS spectrum [2]. Analysis of the genetic overlaps between FTD and ALS has identified multiple biological pathways implicated in both diseases, including RNA regulation, mitochondrial dysfunction, protein homeostasis, autophagy and inflammation [2].

TDP-43-positive, cytoplasmic inclusions in affected neurons are the key neuropathological feature in up to 97% of ALS and ∼50% of FTD patients [3], including those with mutations in FTD-ALS genes such as *TBK1* [4–7]. Under normal conditions, TDP-43 is a nuclear protein capable of shuttling between the nucleus and cytoplasm. During FTD and ALS, TDP-43 becomes increasingly mislocalised to the cytoplasm, eventually forming inclusions. TDP-43-positive inclusions have been identified in spinal cord motor neurons in ALS patients as well as the frontal and temporal lobes of FTD patients, suggesting they are closely linked to the neurodegeneration that occurs in these areas [8–10].

TBK1 is a serine-threonine kinase that plays a role in two key pathways associated with FTD/ALS pathogenesis: autophagy and inflammation [11]. *TBK1* mutations have been identified in 1-2% of FTD/ALS patients [12]. Genetic studies done to date have identified nonsense, frameshift, missense and deletion mutations throughout the *TBK1* sequence in both familial and sporadic cases of ALS, FTD and FTD-ALS. Nonsense and frameshift mutations strongly increase the risk of FTD/ALS (odds ratio ∼11.8; [12]) and cause obvious disruption to TBK1 structure, potentially decreasing both TBK1 mRNA and protein expression. This implies that *TBK1* haploinsufficiency contributes to FTD/ALS in these cases. Currently, >50 missense or single amino acid deletion variants in *TBK1* have been reported in FTD or ALS patients [11]. The pathogenicity of these variants is unclear as they are less likely to cause major disruption to TBK1 structure and expression. However, missense and single amino-acid deletions are still associated with an increase in susceptibility to FTD/ALS (odds ratio ∼1.6; [12]), implying that at least some of these variants have a biological effect that contributes to development of disease.

Here we report the functional consequences of 16 *TBK1* missense variants of unknown significance (VUS) with established pathogenic mutations as comparators. Cells transfected with *TBK1* variants were assayed for their effect on TBK1 protein expression and two key TBK1 functions: activation of NF-κB and OPTN binding. Expression of several *TBK1* missense VUS had similar effects to TBK1 loss-of-function mutations, demonstrating decreased NF-κB activation and loss of OPTN binding. The potential pathogenic effect of *TBK1* VUS was also assessed via TDP-43 mislocalisation. *TBK1* missense VUS increased the cytoplasmic/nuclear ratio of TDP-43 and this was found to inversely correlate with OPTN binding. This study confirms the relative importance of the interaction between TBK1 and OPTN, in comparison with TBK1’s kinase activity, to FTD/ALS pathogenesis.

## Methods

### Selection of TBK1 variants

*TBK1* variants of unknown significance previously reported in FTD and/or ALS patients were selected from ClinVar (https://www.ncbi.nlm.nih.gov/clinvar/), AD & FTD Mutation Database (http://www.molgen.vib-ua.be/ADMutations/), and literature search (**Supplementary Table 1**). Sixteen variants were selected along the length of the TBK1 protein, representing each of its functional domains (**Figure 1**). Each variant had a minor allele frequency (MAF) < 0.0001 in gnomAD non-neurological non-Finnish European (NFE) population (https://gnomad.broadinstitute.org/) [13] and was predicted damaging/deleterious by at least one of PolyPhen2 HumVar (http://genetics.bwh.harvard.edu/pph2/) [14] and SIFT (https://sift.bii.a-star.edu.sg/) [15] algorithms. We also examined: two known pathogenic mutants p.E696K [4,16] and p.R440X [4,7,17] (generated by deleting TBK1 codons 440-729); an artificial deletion of the last 40 amino acids of TBK1 (del690-729), disrupting the OPTN-binding site (677-729aa) [18]; and p.V464A, which is a common missense variant (allele frequency in gnomAD non-neurological NFE population = 0.0136) and thus unlikely to be pathogenic.

**Figure 1.**
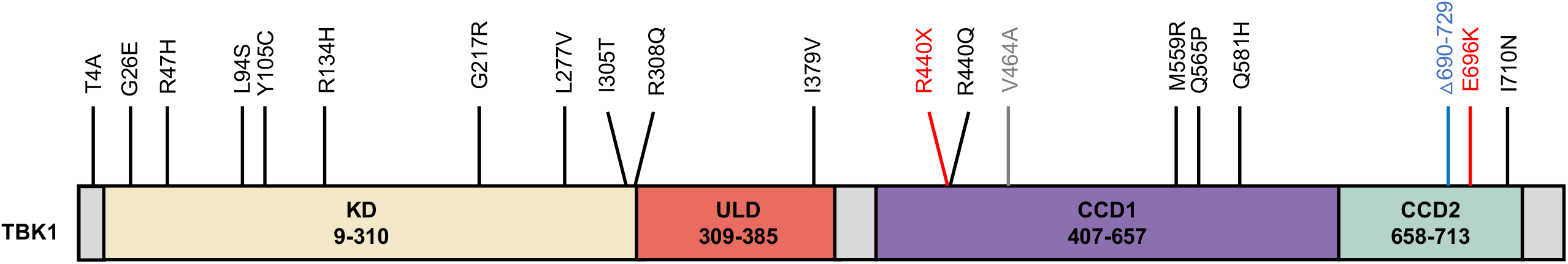
Location of TBK1 variants. Missense variants are indicated in black, known pathogenic variants are indicated in red, the artificial deletion is indicated in blue and the common single nucleotide polymorphism (SNP) in grey. Domains of TBK1 according to UniProt (https://www.uniprot.org/uniprot/Q9UHD2) are indicated. KD = kinase domain, ULD = ubiquitin-like domain, CCD1 = coiled-coil domain 1 and CCD2 = coiled-coil domain 2. The CCD1 domain is also referred to as the scaffold dimerisation domain = SDD [23].

### DNA constructs

*TBK1* mutations were introduced into the pCMV6-TBK1 construct (Origene) by site-directed mutagenesis using the QuikChange Lightning Site-Directed Mutagenesis Kit (Agilent). Mutated *TBK1* cDNA sequences were then subcloned, using *Asi*SI and *Mlu*I restriction sites, into the pCMV6-AC and pCMV6-Entry vectors (Origene) for expression of untagged or C-terminal myc-FLAG-tagged TBK1 protein, respectively. The pCMV6-OPTN vector (Origene) expressing C-terminal myc-FLAG-tagged OPTN protein was used for co-immunoprecipitation experiments. All clones were verified by restriction digestion and sequence analysis.

### Cell Culture

HEK293 and SH-SY5Y were purchased from ATCC and authenticated by STR analysis prior to use in this study. HEK293 cells were maintained in Eagle’s Minimum Essential Medium (EMEM; Gibco) and SH-SY5Y cells in Dulbecco’s Modified Eagle’s Medium/Nutrient Mixture F-12 (DMEM/F12; 1:1 mixture; Gibco) each containing 10% heat-inactivated fetal calf serum (Sigma-Aldrich). For immunocytochemistry, 8-well chamber slides (Ibidi) were coated with poly-L-lysine (Sigma-Aldrich) for 1 h and then washed twice with Dulbecco’s phosphate-buffered saline (DPBS; Gibco).

### Immunoblotting

HEK293 cells were seeded at 1.5 × 10^6^ cells/well in 6-well plates. After 24 h, cells were transfected with untagged wild type or mutant TBK1 constructs (2 μg/well) using Lipofectamine 3000 (6 µL/well; Invitrogen) as per the manufacturer’s protocol. 24 h after transfection, cells were washed with DPBS and lysed by freeze-thawing. 10 µg of protein was resolved by SDS-PAGE on a 4-20% polyacrylamide stain-free gel (Bio-Rad). SDS-PAGE-separated proteins were transferred on to PVDF membrane before blocking and incubation with rabbit anti-TBK1 (1:2000; Abcam ab40676; 1:600; Abcam ab186470) or mouse anti beta-actin (1:10000; Merck #MAB1501) primary antibody in overnight at 4°C. Proteins were visualised by incubation with horseradish peroxidase-conjugated goat anti-rabbit secondary antibody (1:10000; ThermoFisher Scientific) or goat anti-mouse secondary antibody (1:10000; ThermoFisher Scientific) followed by enhanced chemiluminescence using SuperSignal West Pico PLUS Chemiluminescent Substrate (ThermoFisher Scientific) and the ChemiDoc MP Imaging System (Bio-Rad). TBK1 protein expression was normalised to total protein using ImageLab (Bio-Rad).

### NF-κB Luciferase Assay

NF-κB activity was assayed by luciferase reporter assay using the pGL4.32 construct (Promega), which expresses firefly luciferase protein under the control of a NF-κB responsive promoter, and normalizing to Renilla luciferase expression from the pRL-TK construct (Promega), as previously described [19]. Cells were co-transfected with 6.5 ng pGL4.32, 9.75 ng pRL-TK, and 40 ng untagged TBK1 wild type or mutant cDNA construct per well using Lipofectamine 2000 (Invitrogen). Twenty-four hours after transfection, cells were washed with PBS and luciferase expression was assayed using the Dual-Luciferase Reporter Assay System (Promega) and the CLARIOStar microplate reader (BMG Labtech), according to manufacturers’ directions. For each construct combination, the ratio of NF-κB-driven firefly luciferase activity to Renilla luciferase activity was calculated per well and mean firefly: Renilla luciferase ratio was calculated across six replicate wells.

### Co-immunoprecipitation

HEK293 cells were seeded in duplicates at 1.2 × 10^6^ cells/well in 6-well plates. Cells were transfected after 24 h, using Lipofectamine 3000 (Invitrogen) according to manufacturer’s directions. Cells were co-transfected with wild type or mutant TBK1-expressing cDNA constructs plus OPTN-myc-FLAG or empty myc-FLAG vector pCMV6-Entry (2 μg/construct/well). Twenty-four hours after transfection, cells were harvested by EDTA treatment and pelleted by centrifugation. Myc-FLAG-tagged proteins and their interactants were purified from cell pellets using the FLAG Immunoprecipitation Kit (Sigma-Aldrich) and Pierce spin columns (Thermo Fisher Scientific), according to the manufacturers’ directions. Samples were eluted in 40 µL FLAG elution buffer (150 ng/µL 3X FLAG peptide, 50 mM Tris HCl pH 7.5, 150 mM NaCl). 4 µL of lysate and 16 µL of eluate were resolved by SDS-PAGE and immunoblotting was performed as described above using rabbit anti-TBK1, mouse anti-myc (1:2000; Merck 05-724) or mouse anti beta-actin primary antibodies. TBK1 protein expression was normalised to myc (OPTN) expression using ImageLab (Bio-Rad).

### Immunocytochemistry

SH-SY5Y cells were seeded at 7.5 × 10^4^ cells/well and HEK293 cells were seeded at 8 × 10^4^ cells/well. Cells were transfected with TBK1-myc-FLAG constructs (250 ng/well) for 24 (HEK293) or 48 (SH-SY5Y) hours using Lipofectamine 3000 (0.75 µL/well; Invitrogen) as per the manufacturer’s protocol. Transfected cells were fixed and stained as previously described [20] with primary antibodies: rabbit anti-TDP-43 (1:500; Proteintech #10782-2-AP) and mouse anti-FLAG (1:500; Sigma-Aldrich #F1804) antibodies. The next day, cells were incubated with secondary antibodies: rabbit Alexa Fluor-647 (1:500; ThermoFisher Scientific #A-21244) and mouse Alexa Fluor-488 (1:500; ThermoFisher Scientific #A11001), followed by mounting with DAKO fluorescent mounting medium containing 4’,6-diamidino-2-phenylindole (DAPI; Agilent) as previously described [20]. Cells were imaged for fluorescence intensity quantification with a 40x objective on a Nikon A1R confocal microscope. Images of representative cells were obtained with a 100x objective for figure preparation. Laser intensity, gain and offset settings were kept constant and at least ten random fields of view were imaged for each experiment.

### Fluorescence Intensity Quantification

Cytoplasmic mislocalisation of endogenous TDP-43 was quantified using CellProfiler 3.1.9 software (Broad Institute, Cambridge, MA) [21], as previously described [22]. Briefly, nuclei were identified via analysis of the DAPI channel using the *Identify Primary Objects* module and a global, three-class Otsu thresholding method. Strict filtering for cell diameter (30-60 pixels) eliminated any irregular or overlapped nuclei that were present. TDP-positive cells were associated with their corresponding nuclei, using the *Identify Secondary Objects* module and cell area filtering (maximum 3000 pixels) to remove any clumped cells. The *Relate* and *Filter* modules were used to link filtered nuclei to TDP-43-positive cells. Centroid distance filtering (maximum 12 pixels) was applied to remove background fluorescence. FLAG (TBK1)-positive cells were also identified using the *Identify Primary Objects* module. The *Relate* and *Filter* modules were used to select only TDP-43-and FLAG (TBK1)-positive cells and link them to their corresponding nuclei. In each TDP-43-positive cell, the nucleus and cytoplasm were separated into individual objects using the *Identify Tertiary Objects* module. Integrated (total) fluorescence intensity for both cellular compartments was then measured from the TDP-43 channel image using the *Measure Object Intensity* module. Using these measurements, the cytoplasmic to nuclear ratio of TDP-43 was calculated for 250-300 cells per group, for each of the three replicates of the experiment. All filtering and thresholding steps in each experimental pipeline were kept consistent throughout all experimental replicates.

### Statistical Analysis

Data are presented as mean ± standard error of the mean (SEM) from at least three independent experiments. Quantification of immunoblots, NF-κB activity and TDP-43 cytoplasmic to nuclear ratio was normalised to TBK1 WT and compared by repeated measures one-way ANOVA and Dunnett’s multiple comparisons test. Pearson correlation was used to analyse the relationship between the results of each assay. All statistical analyses were performed using GraphPad Prism 9. Significance for all tests was set at p < 0.05.

## Results

### Analysis of steady-state TBK1 protein levels

We selected 16 missense VUS for analysis (**Supplementary Table 1**) that were distributed along the TBK1 protein, representing all of its functional domains (**Fig. 1**).

We performed immunoblot analysis of TBK1 protein expression in HEK293 cells transiently transfected with wild-type or variant TBK1 cDNA constructs (**Supplementary Fig. 1**). TBK1_G217R_ protein was present at significantly lower levels relative to TBK_WT_ (0.40±0.03, p=0.0123), in line with previous results [24]. Conversely, TBK1_Q565P_ was expressed at significantly higher levels relative to wild-type (2.03±0.09, p=0.0329). No significant differences in steady-state TBK1 levels were observed for other TBK1 variants.

### TBK1 missense variants differentially alter NF-κB activation

As a serine-threonine kinase belonging to the noncanonical IκB kinase family, TBK1 plays a key regulatory role in many pathways via phosphorylation of its substrates, including the NF-κB complex [25,26]. We investigated the impact of TBK1 missense VUS on NF-κB activation by luciferase reporter assay in HEK293 cells overexpressing wild-type or variant TBK1 (**Fig. 2**) as a readout of how these variants affect TBK1 kinase activity. As expected, transfection with TBK1_WT_ led to a significant increase in NF-κB activity relative to empty vector (0.34±0.03, p<0.0001). For four missense variants, all located in the kinase domain, a significant reduction in NF-κB activity was observed relative to wild-type: TBK1_G26E_ (0.56±0.05, p=0.0043), TBK1_L94S_ (0.44±0.03, p=0.0003), TBK1_R134H_ (0.44±0.03, p=0.0003) and TBK1_G217R_ (0.42±0.05, p=0.0016). Nonsense mutant TBK1_R440X_ showed a similar reduction relative to wild-type (0.33±0.06, p=0.0017). Conversely, a significant increase in NF-κB activation, albeit of modest effect, was observed for TBK1_I379V_ located in the ubiquitin-like domain (1.29±0.04, p=0.0094). Large increases in NF-kB activation were observed for coiled-coil domain 1 (CCD1) variants TBK1_R440Q_ (6.42±0.75, p=0.0089) and TBK1_Q565P_ (9.16±0.98, p=0.0052); and significant increases were also observed for pathogenic mutant TBK1_E696K_ (1.79±0.16, p=0.0294) and the TBK1_Δ690-729_ deletion (2.70±0.34, p=0.0332).

**Figure 2.**
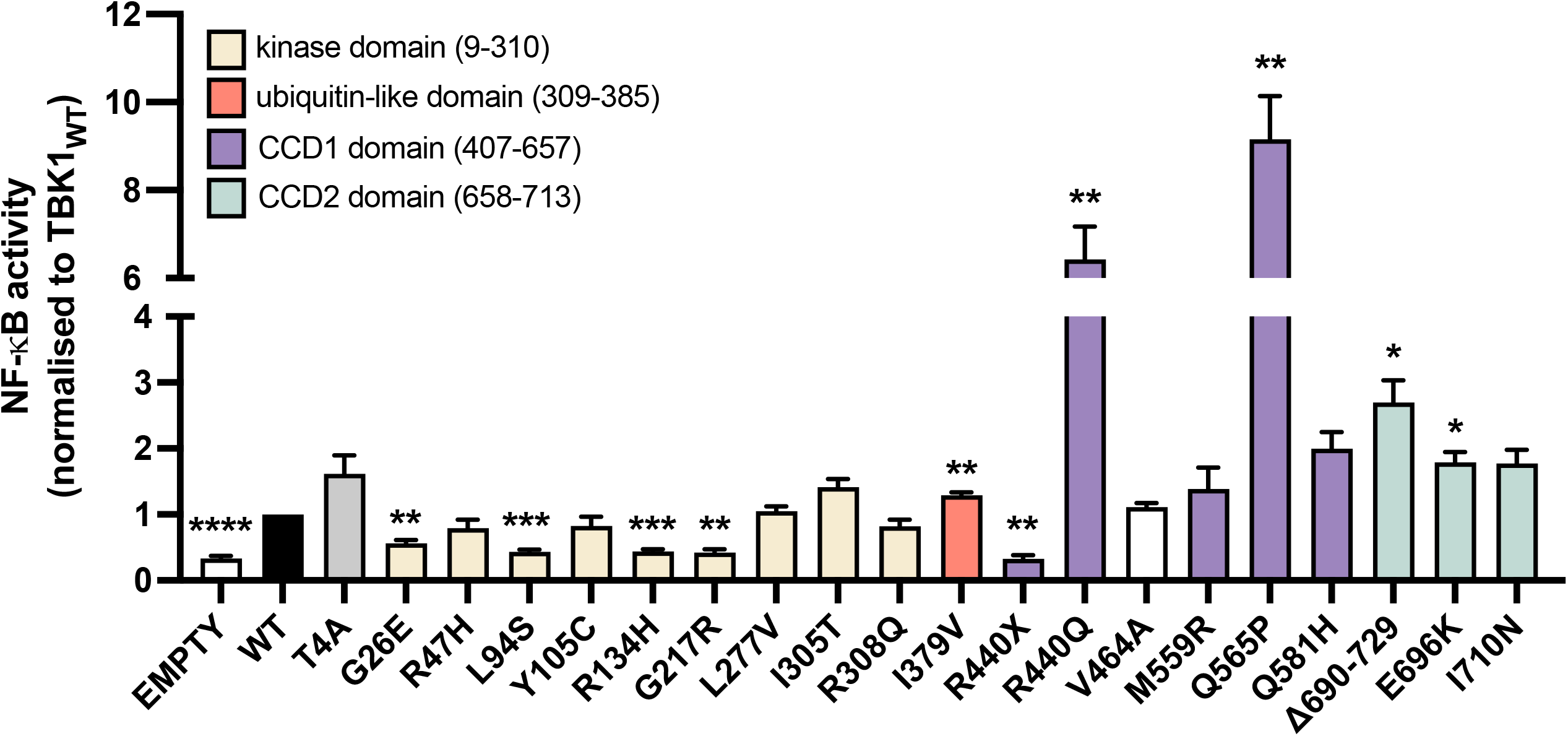
*TBK1* missense variants show differential effects on NF-κB activation. Quantification of NF-κB activity in HEK293 cells transiently transfected with NF-κB-responsive luciferase construct pGL4.32 plus empty vector, TBK1_WT_ or TBK1 variant constructs (n=5). TBK1_WT_ led to a significant increase in NF-κB activity relative to empty vector (0.34±0.03, p<0.0001). TBK1_G26E_ (0.56±0.05, p=0.0043), TBK1_L94S_ (0.44±0.03, p=0.0003), TBK1_R134H_ (0.44±0.03, p=0.0003), TBK1_G217R_ (0.42±0.05, p=0.0016), TBK1_I379V_ (1.29±0.04, p=0.0094), TBK1_R440X_ (0.33±0.06, p=0.0017), TBK1_R440Q_ (6.42±0.75, p=0.0089), TBK1_Q565P_ (9.16±0.98, p=0.0052), TBK1_Δ690-729_ (2.70±0.34, p=0.0332) and TBK1_E696K_ (1.79±0.16, p=0.0294) all caused significant changes in NF-κB activity relative to TBK1_WT_. NF-κB activity was compared using repeated measures one-way ANOVA and Dunnett’s multiple comparisons test. Data is represented as mean ± SEM. *p<0.05; **p<0.01; ***p<0.001; ****p<0.0001.

### TBK1 missense variants differentially alter binding to OPTN

TBK1 is a major binding partner of the autophagy adaptor and FTD/ALS-associated protein, OPTN, enhancing its ability to link ubiquitinated proteins and damaged mitochondria with autophagosome-associated proteins such as LC3-II [11]. In order to assess whether missense VUS in TBK1 affect its ability to interact with OPTN we co-transfected HEK293 cells with OPTN-mycFLAG construct and wild-type or variant TBK1 constructs, followed by immunoprecipitation with anti-FLAG antibody (**Fig. 3**). As expected, TBK1_R440X_ and TBK1_Δ690-729_ constructs lacking an intact coiled-coil domain 2 (CCD2) domain, required for interaction with OPTN [18], showed a complete abolishment of OPTN binding (both 0.00±0.00, p<0.0001). Known pathogenic mutant TBK1_E696K_ also showed a significant decrease in OPTN binding relative to wild-type (0.02±0.02, p=0.0007), in line with previous observations [18]. A significant decrease in the amount of TBK1 bound to OPTN was found for two missense variants located in the kinase domain (TBK1_L94S_: 0.04±0.04, p=0.0060; TBK1_G217R_: 0.27±0.08, p=0.0403), three located in the CCD1 domain (TBK1_R440Q_: 0.17±0.09, p=0.0359; TBK1_M559R_: 0.05±0.05, p=0.0078; TBK1_Q581H_: 0.20±0.09, p=0.0358) and one in the CCD2 domain (TBK1_I710N_: 0.07±0.05, p=0.0079). A non-significant downward trend was also observed for CCD1 variant TBK1_Q565P_ (0.14±0.14, p=0.0735). Conversely, a significant increase in OPTN binding was observed for TBK1_T4A_ (1.93±0.11, p=0.0355) and non-significant upward trends in binding were observed for kinase domain variants TBK1_R47H_ (3.23±0.37, p=0.0726) and TBK1_R134H_ (2.53±0.26, p=0.0743).

**Figure 3.**
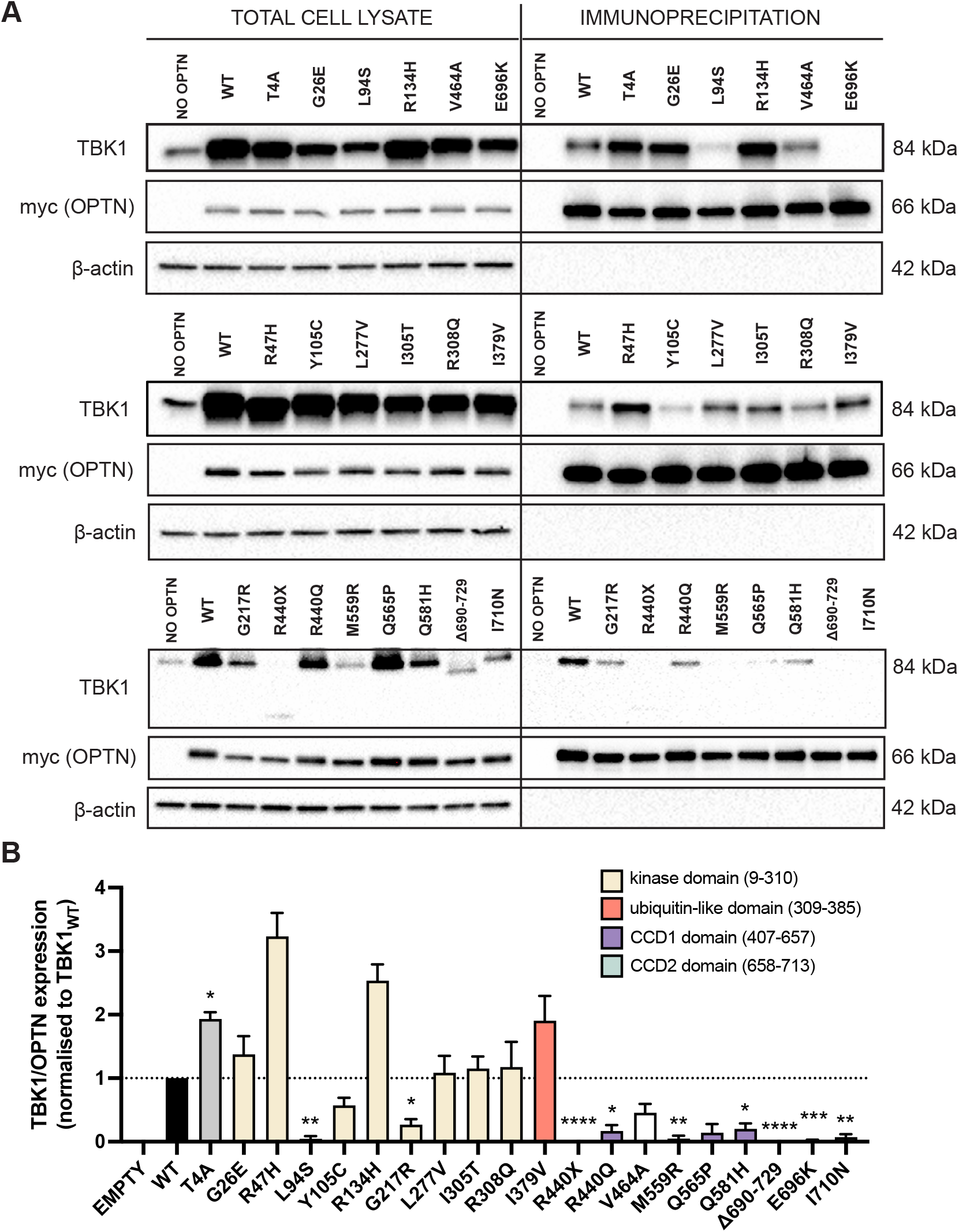
*TBK1* missense variants affect the binding of TBK1 to OPTN. (**a**) Representative immunoblots of HEK293 cell extracts transiently co-transfected with empty-mycFLAG or OPTN-mycFLAG constructs and untagged *TBK1* wild-type (WT) or variant constructs were immunoprecipitated with FLAG antibody. Immunoblots were probed with anti-TBK1, anti-myc and anti-β-actin primary antibodies. (**b**) Quantification of immunoprecipitation immunoblots showing a significant difference in the amount of TBK1 bound to OPTN in a number of TBK1 variants when compared to TBK1_WT_ (n=3). A significant decrease in the amount of TBK1 bound to OPTN was found for TBK1_L94S_: 0.04±0.04, p=0.0060; TBK1_G217R_: 0.27±0.08, p=0.0403; TBK1_R440X_: 0.00±0.00, p<0.0001; TBK1_R440Q_: 0.17±0.09, p=0.0359; TBK1_M559R_: 0.05±0.05, p=0.0078; TBK1_Q581H_: 0.20±0.09, p=0.0358; TBK1_Δ690-729_: 0.00±0.00, p<0.0001; TBK1_E696K_: 0.02±0.02, p=0.0007 and TBK1_I710N_: 0.07±0.05, p=0.0079. A significant increase in OPTN binding was observed for TBK1_T4A_:1.93±0.11, p=0.0355. Bound TBK1 was compared using repeated measures one-way ANOVA and Dunnett’s multiple comparisons test. Data is represented as mean ± SEM. *p<0.05; **p<0.01; ***p<0.001, ****p<0.0001.

### TBK1 missense variants can cause cytoplasmic mislocalisation of TDP-43

We assessed the ability of TBK1 missense variants to cause cytoplasmic mislocalisation of TDP-43, a pathological characteristic of FTD and ALS affected neurons [8–10]. HEK293 cells were transiently transfected with TBK1_WT_-mycFLAG or TBK1 variant-mycFLAG constructs and changes in endogenous TDP-43 subcellular localisation were quantified using indirect immunofluorescence (**Fig. 4, Supplementary Fig. 4-5**). Cells expressing known pathogenic mutants TBK1_R440X_ (**Fig. 4d**) and TBK1_E696K_ (**Fig. 4h**) showed a significant increase in the ratio of cytoplasmic/nuclear TDP-43 relative to those expressing TBK1_WT_ (TBK1_R440X_: 1.19±0.00, p=0.0052; TBK1_E696K_: 1.44±0.00, p=0.0002). We also observed a significant increase in the TDP-43 cytoplasmic/nuclear ratio in cells expressing kinase domain variants TBK1_Y105C_ (1.39±0.03, p=0.0186; **Fig. 4b**) and TBK1_R308Q_ (1.27±0.02, p=0.0218; **Fig. 4c**), CCD1 domain variant TBK1_R440Q_ (1.40±0.05, p=0.00459; **Fig. 4e**) and CCD2 domain variant TBK1_I710N_ (1.23±0.02, p=0.0185; **Fig. 4i**), as well as for CCD2-truncated TBK1_Δ690-729_ (1.38±0.01, p=0.0029; **Fig. 4g**).

**Figure 4.**
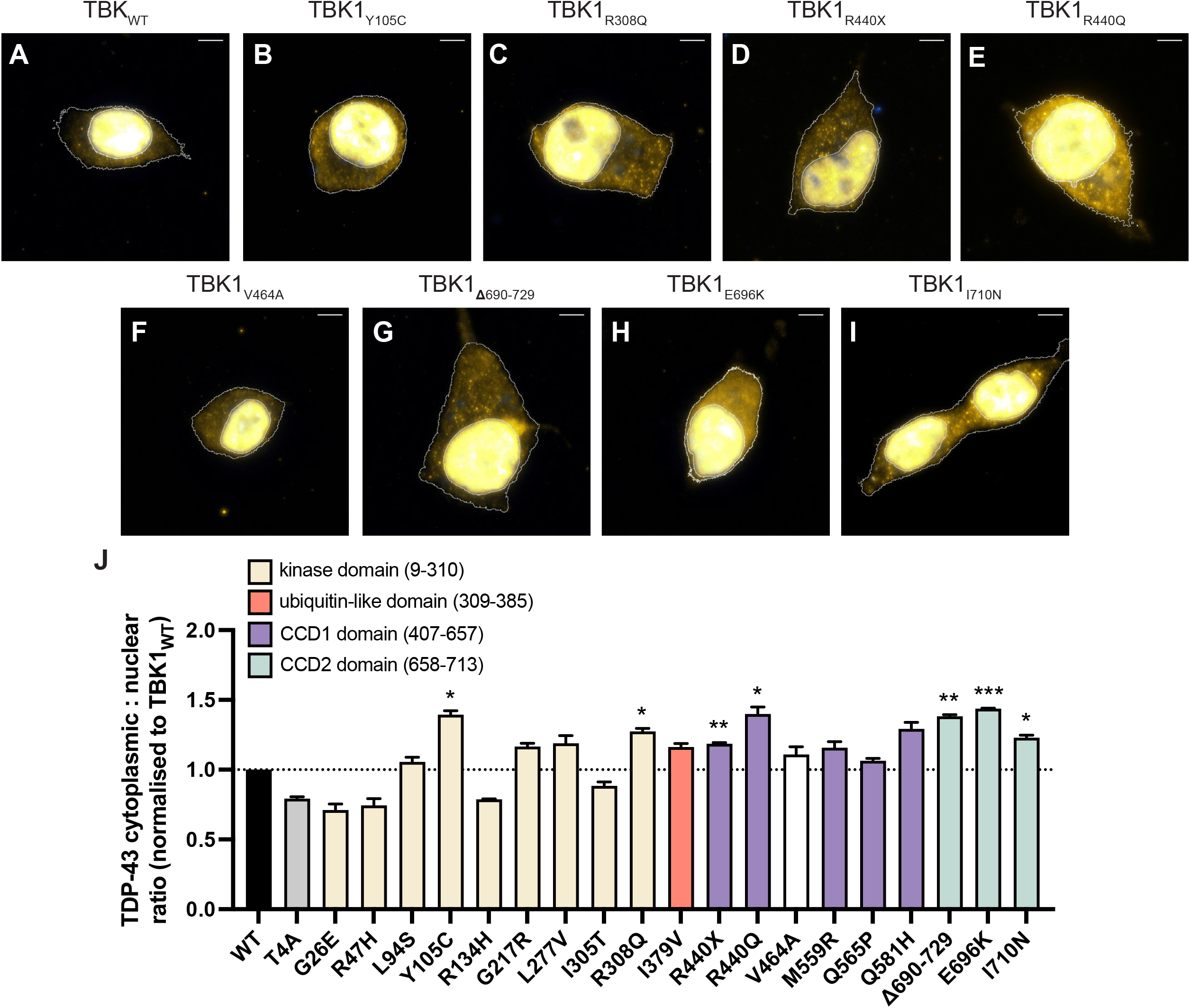
Expression of *TBK1* missense variants induces cytoplasmic mislocalisation of TDP-43. Representative fluorescence images of HEK293 cells expressing mycFLAG-tagged (**a**) TBK1_WT_, (**b**) TBK1_Y105C_ (1.39±0.03, p=0.0186), (**c**) TBK1_R308Q_ (1.27±0.02, p=0.0218), (**d**) TBK1_R440X_ (1.19±0.00, p=0.0052), (**e**) TBK1_R440Q_ (1.40±0.05, p=0.00459), (**f**) TBK1_V464A_, (**g**) TBK1_Δ690-729_, (1.38±0.01, p=0.0029), (**h**) TBK1_E696K_ (1.44±0.00, p=0.0002) or (**i**) TBK1_I710N_ (1.23±0.02, p=0.0185). Images displayed are merged DAPI-stained nuclei (blue) and TDP-43 (yellow) channels. Nuclei and cell membrane outlines are defined by DAPI and TBK1 (FLAG) staining, respectively. Scale bars = 5 µm. (**j**) Quantification of the cytoplasmic/nuclear ratio of endogenous TDP-43 shows an increase in cells expressing some TBK1 missense variants, known pathogenic mutations and deletion constructs (n=3 biological replicates of 300 cells each). Ratios were compared using repeated measures one-way ANOVA and Dunnett’s multiple comparisons test. Data is represented as mean ± SEM. *p<0.05; **p,0.01; ***p<0.001.

In order to confirm whether the effect of TBK1 variants on TDP-43 mislocalisation was also observed in neuronal-like cells, we repeated this experiment in SH-SY5Y neuroblastoma cells for selected TBK1 variants with a normalised cytoplasmic/nuclear ratio of TDP-43 >1 (**Supplementary Fig. 6**). All significant effects in HEK293 cells were replicated in SH-SY5Y cells, and additionally the cytoplasmic mislocalisation of TDP-43 in SH-SY5Y cells was statistically significant for variants TBK1_G217R_ (1.27±0.00, p=0.0002) and TBK1_Q581H_ (1.10±0.00, p=0.005; **Supplementary Fig. 6d, k, o**).

### The degree of TBK1 and OPTN binding inversely correlates with TDP-43 cytoplasmic mislocalisation

To establish the relationships between each of the TBK1 functions we had previously assessed we performed correlation analysis between assay data for each of the variants (**Fig. 5a, Supplementary Fig. 7**). TBK1 protein levels were significantly positively correlated with NF-κB activation (r = 0.621; p=0.0035; **Fig. 5b**), however this was no longer significant after removal of one outlier (r = 0.043, p = 0.8612; **Supplementary Fig. 7a**). The degree of OPTN binding relative to wild-type TBK1 showed a significant inverse correlation with degree of cytoplasmic mislocalisation of TDP-43 (r = -0.706; p=0.0005, **Fig. 5c**). Neither TBK1 protein levels nor NF-kB activation were significantly correlated with TDP-43 mislocalisation (**Supplementary Fig. 7c, e**)

**Figure 5.**
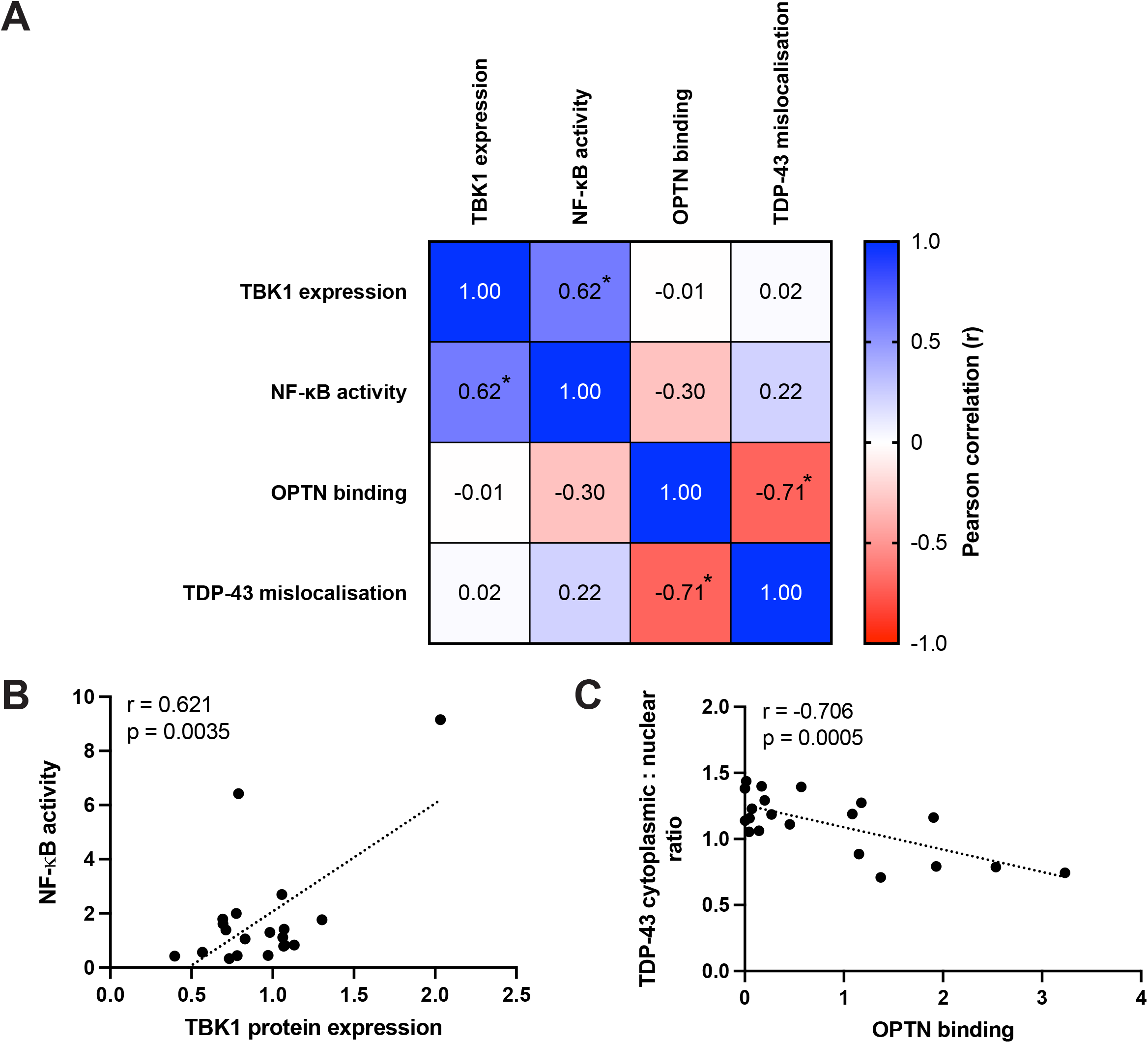
Correlation analysis of TBK1 functional assays. (**a**) Pearson correlation matrix of TBK1 functional assays performed. Labels indicate Pearson correlation coefficient (r) from analysis and asterisks indicate significant correlations. (**b**) Significant correlation between TBK1 protein expression and NF-κB activity of TBK1 variants. (**c**) Significant inverse correlation between amount of TBK1 bound to OPTN and the TDP-43 cytoplasmic to nuclear ratio of TBK1 variants. r = Pearson correlation coefficient.

## Discussion

To date, greater than 90 *TBK1* variants have been identified in patients with ALS, FTD or FTD-ALS, the majority of which are missense VUS [11]. This study sought to identify regions and/or functions of TBK1 that were involved in the pathogenesis of FTD/ALS through a battery of cellular based assays. Through co-immunoprecipitation experiments we discovered that expression of many *TBK1* VUS, particularly in the CCD1 and CCD2 domains of TBK1 decreased OPTN binding efficiency. These results also inversely correlated with TDP-43 mislocalisation. These findings indicate that disruption of the interaction between TBK1 and OPTN may be involved in FTD/ALS pathogenesis.

TBK1 is a multifunctional kinase that plays a regulatory role in many signalling pathways, via phosphorylation of its substrates. During NF-κB activation, TBK1 phosphorylates multiple members and effectors in the NF-κB signalling pathway [27], therefore detection of NF-κB activation can be viewed as a non-substrate-specific assay of TBK1 kinase activity. As expected, the four missense VUS that led to a significant reduction in NF-κB activity (TBK1_G26E_, TBK1_L94S_, TBK1_R134H_ and TBK1_G217R_) were all located in TBK1’s kinase domain (aa9-310). The nonsense mutant TBK1_R440X_ also showed a similar reduction in NF-κB activation. The results for TBK1_L94S_ TBK1_R134H_ and TBK1_G217R_ are broadly in line with data from previous studies [24,28]. The reduced TBK1 protein levels shown with expression of TBK1_G217R_ in previous studies [24,29,30], and its segregation with disease in multiple affected family members [29] have led to TBK1_G217R_ being classified as likely pathogenic. We observed no significant change in kinase activity in TBK1_R47H_ and TBK1_M559R_ cells, which showed reduced NF-κB activation in a previous study [24]. This study measured NF-κB activity using a different reporter system, and in a TBK1-knockout cell line, which may account for the discrepancy between our findings and theirs. It is also not clear whether biological replicates were performed.

We identified three *TBK1* VUS (TBK1_I379V_, TBK1_R440Q_ and TBK1_Q565P_) as well as the pathogenic mutation TBK1_E696K_ and the artificial deletion (TBK1_Δ690-729_), outside the kinase domain that significantly increased TBK1 kinase activity, two of which have previously demonstrated comparable kinase activity to TBK1_WT_ (TBK1_Q565P_ and TBK1_E696K_) [24]. Whilst the increase in NF-κB activation shown in TBK1_Q565P_ cells is likely due to increased TBK1 expression (**Supplementary Fig. 1**), the reason for increased kinase activity with other TBK1 variants is unknown. The marked increase in NF-κB activation demonstrated by TBK1_Q565P_ drives the significant correlation between TBK1 protein expression and NF-κB activity (**Fig. 5b, Supplementary Fig. 7a**). Increased TBK1 protein expression and kinase activity have previously been associated with obesity [31,32] and *TBK1* gene duplication is associated with glaucoma [33,34]. However, in FTD/ALS, TBK1 is typically associated with haploinsufficiency and therefore decreased protein expression and function and it is not clear whether increased TBK1 kinase activity is of clinical relevance to FTD/ALS pathogenesis.

TBK1 directly interacts with OPTN, an autophagy adaptor protein that links ubiquitinated proteins and damaged mitochondria with autophagosome-associated proteins such as LC3-II [11]. The interaction between TBK1 and OPTN occurs via the CCD2 domain of TBK1 (aa658-713) [18]. Investigation of this interaction showed that several *TBK1* VUS located in the CCD1 domain caused a significant decrease in OPTN binding, indicating that structural integrity of this domain may also be important to the TBK1-OPTN interaction. TBK1_Q565P_ showed a non-significant downward trend in OPTN binding similar to a previous study [24]. TBK1_E696K_, TBK1_M559R_ and deletion mutants in the CCD2 domain (p.690-713del) have previously been shown to abolish OPTN binding [4,18,24] and this was also confirmed in our study. Kinase domain variants TBK1_L94S_ and TBK1_G217R_ also showed a reduction in OPTN binding, which for TBK1_G217R_ has been reported previously [24,29]. We did not observe a reduction in OPTN binding for TBK1_R308Q_, in contrast to a previous study, although it is not clear whether biological replicates were performed [24] One TBK1 VUS (TBK1_T4A_) led to a significant increase in OPTN binding. Enhanced binding of TBK1 and OPTN has been reported in cells expressing the familial primary open-angle glaucoma mutation OPTN_E50K_ [35] but it is not clear whether this would be of relevance to FTD or ALS pathogenesis.

Mislocalisation of neuronal TDP-43 from the nucleus to the cytoplasm is a key pathogenic process observed in the majority of ALS and ∼50% of FTD patients [3], including those with *TBK1* mutations [4,36,37]. *TBK1* knockdown human cellular and mouse models have also demonstrated TDP-43 cytoplasmic mislocalisation [38,39]. Here we assess for the first time the effect of *TBK1* missense variants on this process. Importantly, both established pathogenic mutations TBK1_R440X_ & TBK1_E696K_ and the likely pathogenic variant TBK1_G217R_ showed significant TDP-43 mislocalisation in SH-SY5Y cells, validating this approach to assess pathogenicity of TBK1 variants. In HEK293 and SH-SY5Y cells, a higher proportion of CCD1 or CCD2 domain variants (8/8, 100%) showed a significant or trend significant mislocalisation of TDP-43 in comparison with kinase domain variants (2/9, 22%), indicating that *TBK1* VUS in the CCD1 and CCD2 domains are more likely to be pathogenic.

Whilst the >90 *TBK1* variants identified in FTD/ALS patients are spread along the *TBK1* sequence more than half are located in the CCD1 and CCD2 domains, including a higher proportion of deletion, nonsense, splice, frameshift and insertion variants (26/40 variants; 65%) compared to the similarly sized kinase domain (11/36 variants; 31%) [11]. The significant inverse correlation we discovered between OPTN binding and TDP-43 mislocalisation indicates that this relationship may be crucial to FTD/ALS pathogenesis (**Fig. 5c**). Conversely, we saw no significant correlation between TBK1 kinase activity, as measured by NF-κB activation, and TDP-43 mislocalisation (**Supplementary Fig. 7e**), indicating that a reduction in kinase activity may not be pathogenic in the context of FTD & ALS. Decreased binding of OPTN to TBK1 has the potential to reduce the efficiency of OPTN linking ubiquitinated cargo to LC3II and subsequently autophagosome formation and maturation [11,40]. Accumulation of unwanted and/or damaged proteins in cytosolic aggregates (e.g. TDP-43) is characteristic of FTD/ALS and ultimately leads to cell death.

Our study had certain limitations. By using an NF-κB luciferase reporter assay we were able to assess TBK1 kinase activity in a more quantitative and relatively substrate-agnostic manner compared to western blot analysis of substrate phosphorylation. However, we cannot exclude the possibility that reduced phosphorylation of a specific TBK1 substrate may play a role in FTD/ALS. Secondly, we selected OPTN binding for analysis due to the discovery of TBK1 pathogenic variants p.E696K and p.690-713del which abrogate OPTN binding [4], indicating that loss of the TBK1-OPTN interaction may be sufficient to cause disease. Our findings support this conclusion: however, it remains possible that *TBK1* variants causing FTD/ALS may do so via loss of interaction with another protein binding to the same region as OPTN. Lastly, in this study we utilised HEK293 and SH-SY5Y cell lines due to their relatively high transfection efficiency and replicability of results, allowing examination of a large number of variants. Further investigation of the putative pathogenic *TBK1* missense variants identified in this study in lower-throughput but more physiologically relevant cellular and/or animal models is now warranted, in order to further clarify the biological consequences of these variants.

## Conclusions

Overall, this study has shown a significant inverse correlation between the ability of TBK1 to bind OPTN and TDP-43 mislocalisation, one of the pathological hallmarks of FTD/ALS. By testing a selection of *TBK1* VUS spread along the *TBK1* sequence we have been able to isolate the CCD1 and CCD2 domains and the relationship between TBK1 and OPTN as areas of interest for future research. This study also examined for the first time the effect of *TBK1* VUS on a neuropathology-relevant phenotype, namely TDP-43 cytoplasmic mislocalisation, and demonstrated the value of this assay as an initial readout for pathogenicity. This simple *in vitro* assay can now be applied to other disease proteins (e.g. fused in sarcoma (*FUS*)) or to other genes associated with FTD/ALS, in conjunction with functional readouts to help isolate regions and/or functions of importance to disease development.

## Abbreviations

ALS: amyotrophic lateral sclerosis
CCD1: coiled-coil domain 1
CCD2: coiled-coil domain 2
CHCHD10: coiled-coil-helix-coiled-coil-helix domain containing 10
DAPI: 4’,6-diamidino-2-phenylindole
DMEM: Dulbecco’s Modified Eagle’s Medium
DPBS: Dulbecco’s phosphate-buffered saline
EMEM: Eagle’s Minimum Essential Medium
FTD: frontotemporal dementia
FTD-ALS genes: genes with mutations that can lead to either FTD or ALS
FUS: fused in sarcoma
MAF: minor allele frequency
NFE: non-Finnish European
OPTN: optineurin
SEM: standard error of the mean
SNP: single nucleotide polymorphism
SQSTM1: sequestosome 1
TBK1: TANK-binding kinase 1
VCP: valosin containing protein
VUS: variants of unknown significance
WT: wild-type

## Declarations

### Ethics approval and consent to participate

Not applicable.

### Consent for publication

Not applicable.

### Availability of data and materials

The datasets and materials generated during the current study are available from the corresponding author on reasonable request.

### Authors’ contributions

LJO, JBK and CDS conceived the study. LF, LMB and MH generated and sequence-verified mutant cDNA constructs. LMB, LJO and LF carried out the immunoblot and co-immunoprecipitation experiments. MH and JZ carried out the NF-KB experiments and LJO performed the TDP-43 mislocalisation experiments. LJO and CDS participated in data analysis. LJO and CDS drafted the manuscript. All authors read and approved the final manuscript.

## Acknowledgements

The authors acknowledge the facilities and technical assistance of Microscopy Australia at the Australian Centre for Microscopy & Microanalysis at the University of Sydney.

