## Supplementary Figures and Table for "TDP-43 subcellular mislocalisation is correlated with loss of optineurin binding for frontotemporal dementia and amyotrophic lateral sclerosis associated *TBK1* missense variants"

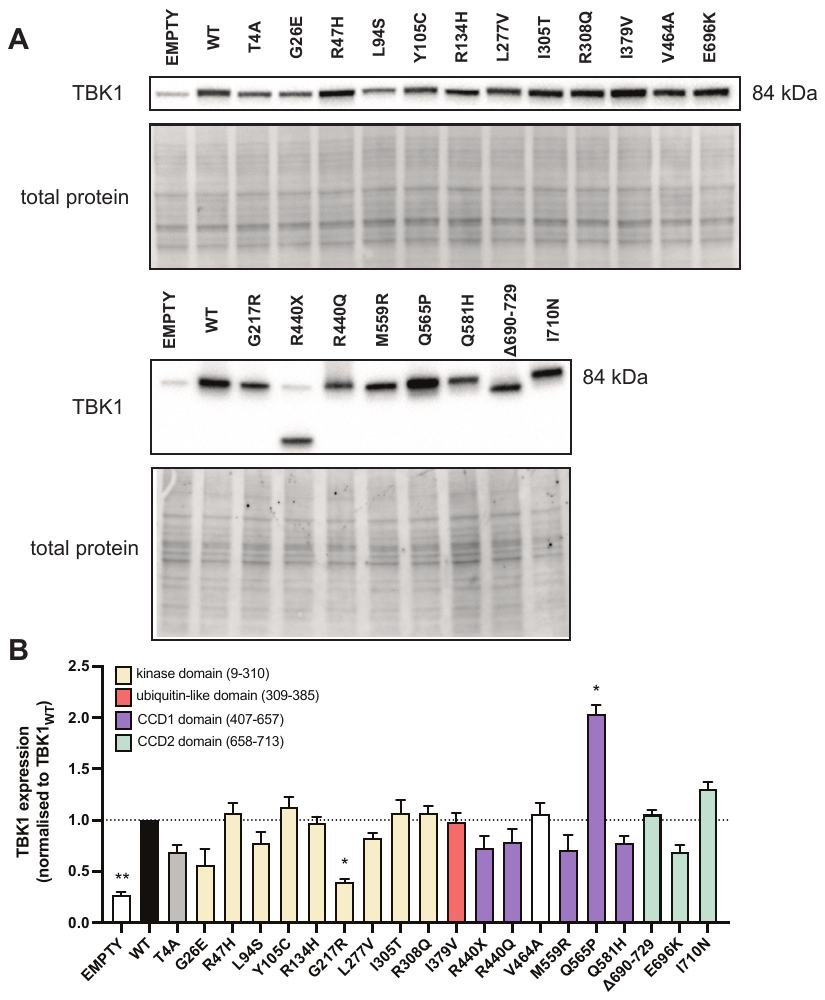

**Supplementary Figure 1.** TBK1 protein expression in HEK293 cells expressing TBK1 variants. (**a**) Representative immunoblots of TBK1 protein expression (**b**) Quantification of TBK1 expression normalised to TBK1_WT_ (n=3). Analysis done using repeated measures one-way ANOVA and Dunnett’s multiple comparisons test. Data is represented as mean ± SEM. *p<0.05; **p<0.01.

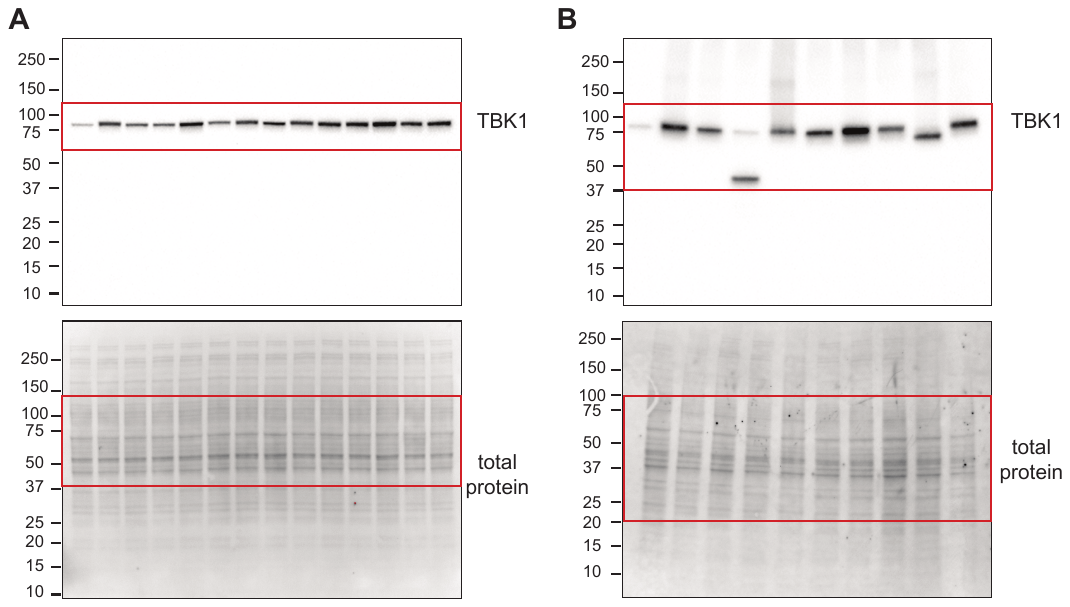

**Supplementary Figure 2.** Uncropped images of immunoblots from Supplementary Figure 1a.

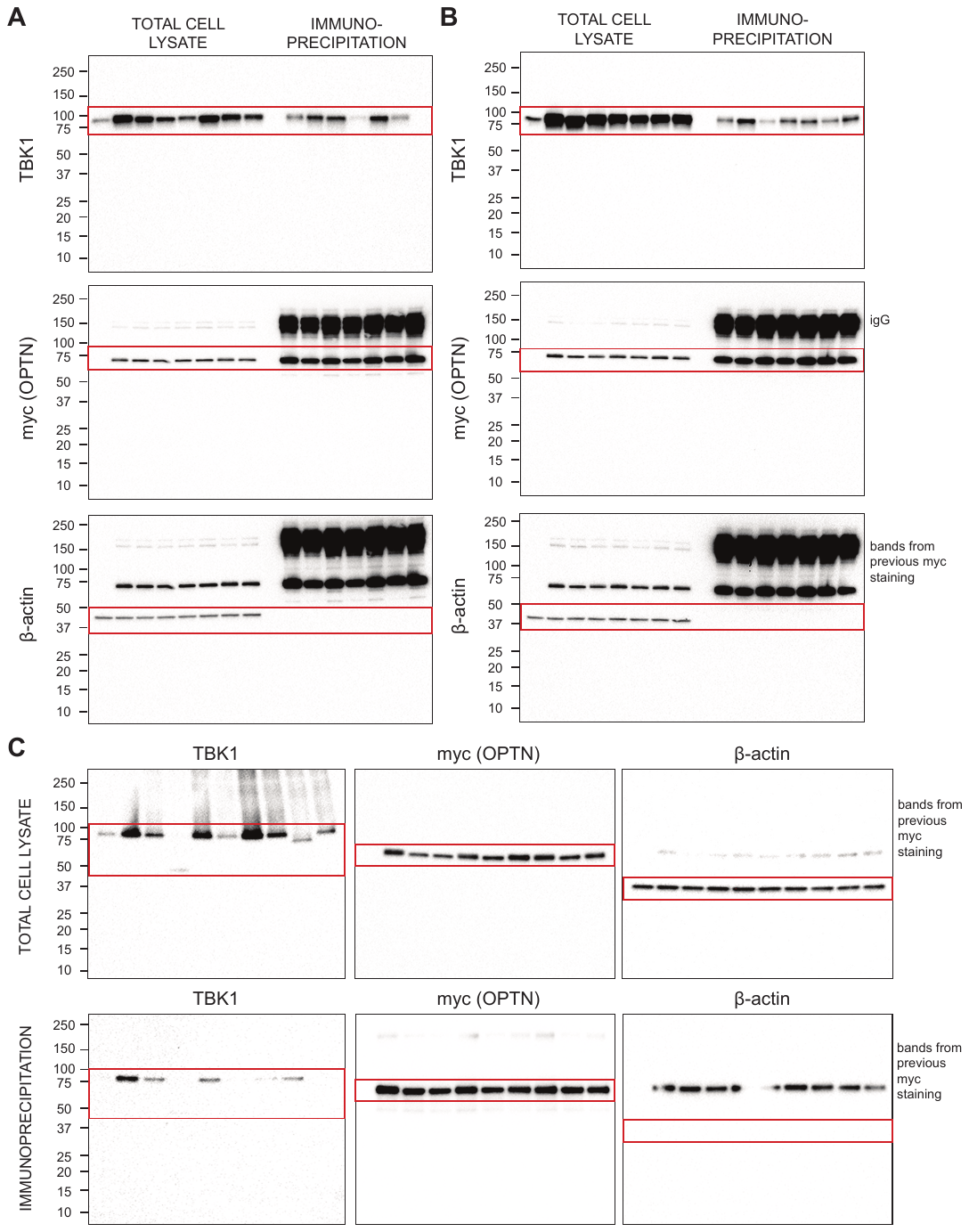

**Supplementary Figure 3.** Uncropped images of immunoblots from Figure 3a.

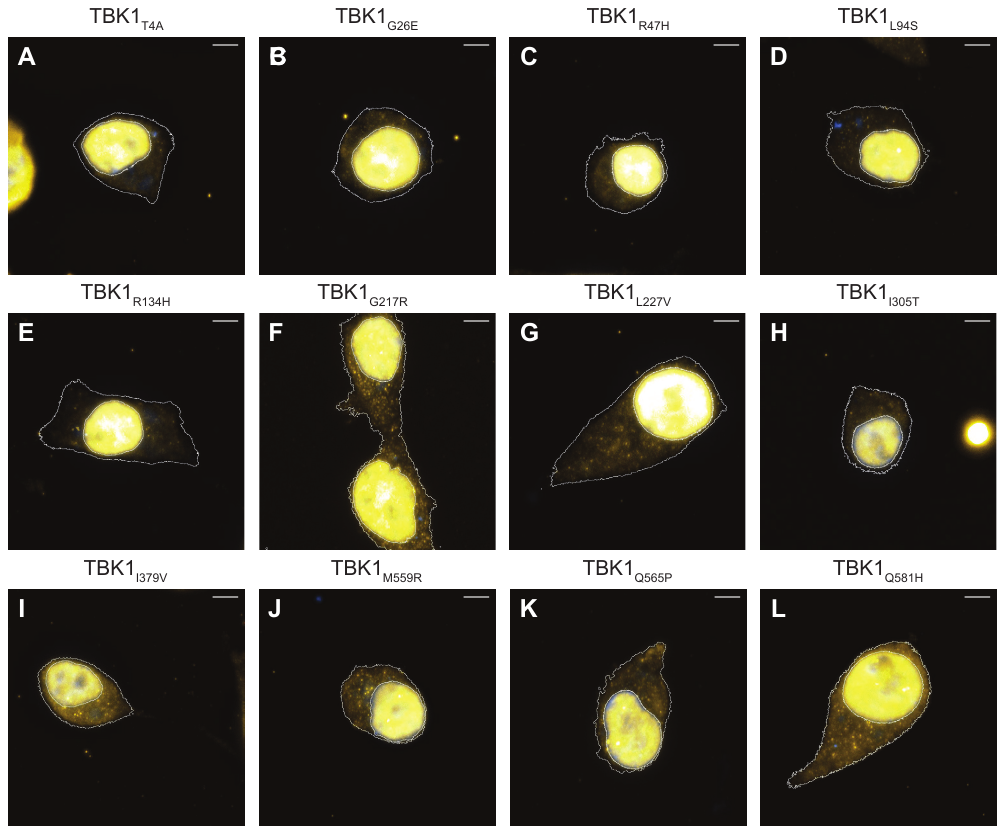

**Supplementary Figure 4.** Representative fluorescence images of HEK293 cells expressing mycFLAG-tagged TBK1 variants. (**a**) TBK1_T4A_, (**b**) TBK1_G26E_, (**c**) TBK1_R47H_, (**d**) TBK1_L94S_, (**e**) TBK1_R134H_, (**f**) TBK1_G217R_, (**g**) TBK1_L227V_, (**h**) TBK1_I305T_, (**i**) TBK1_I379V_, (**j**) TBK1_M559R_, (**k**) TBK1_Q565P_ or (**l**) TBK1_Q581H_. Images displayed are merged DAPI-stained nuclei (blue) and TDP-43 (yellow) channels. Nuclei and cell membrane outlines are defined by DAPI and TBK1 (FLAG) staining, respectively. Scale bars = 5 μm.

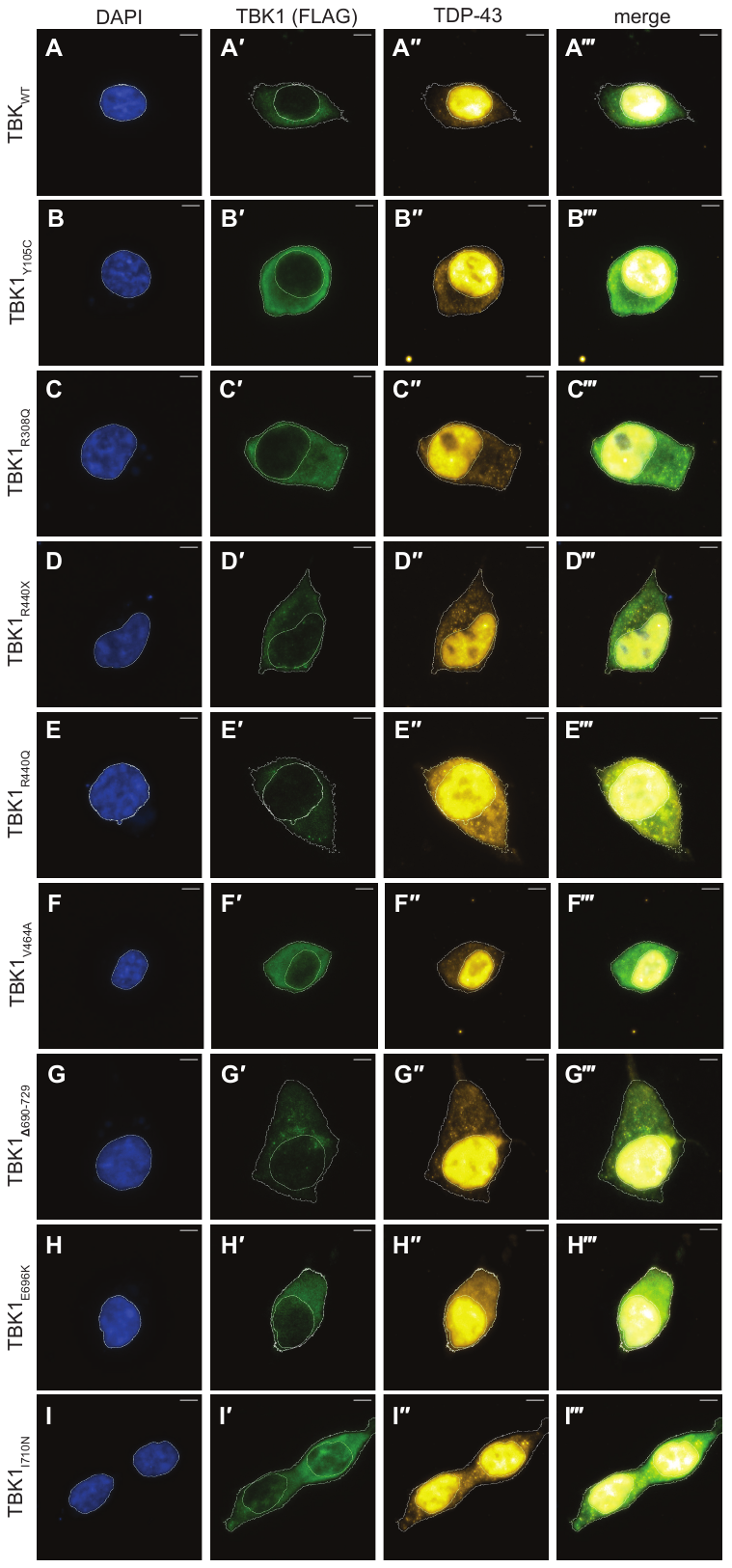

**Supplementary Figure 5.** Expression of TBK1 missense variants induces cytoplasmic mislocalisation of TDP-43. Representative fluorescence images of HEK293 cells expressing mycFLAG-tagged (**a**) TBK1_WT_, (**b**) TBK1_Y105C_, (**c**) TBK1_R308Q_, (**d**) TBK1_R440X_, (**e**) TBK1_R440Q_, (**f**) TBK1_V464A_, (**g**) TBK1_Δ690-729_, (**h**) TBK1_E696K_ or (**i**) TBK1_I710N_. Images displayed are DAPI-stained nuclei (blue), TDP-43 (yellow), TBK1 (FLAG) and merged channels. Nuclei and cell membrane outlines are defined by DAPI and TBK1 (FLAG), respectively. Scale bars = 5 μm.

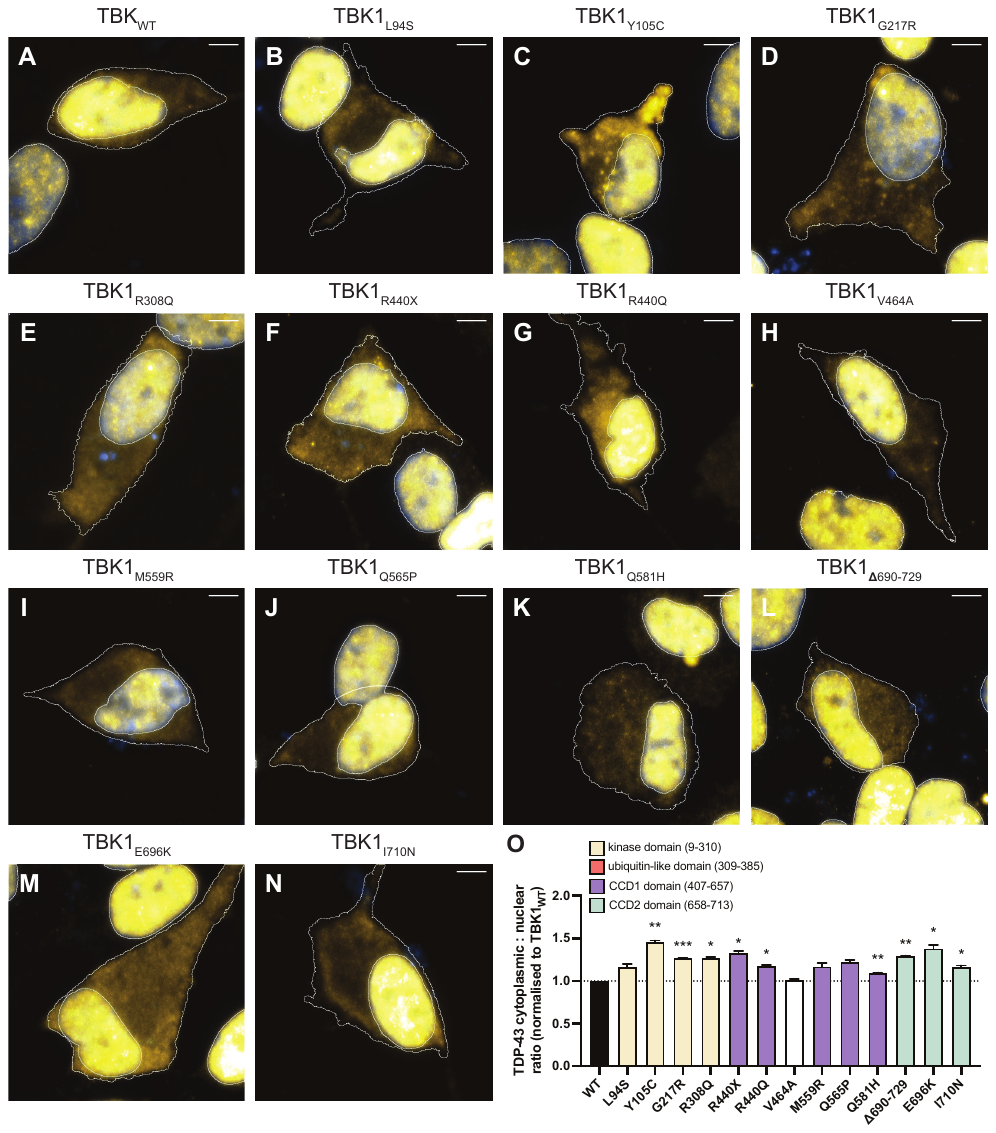

**Supplementary Figure 6.** Expression of TBK1 missense variants induces cytoplasmic mislocalisation of TDP-43 in SH-SY5Y cells. Representative fluorescence images of SH-SY5Y cells expressing mycFLAG-tagged (**a**) TBK1_WT_, (**b**) TBK1_L94S,_ (**c**) TBK1_Y105C_ (1.46±0.02, p=0.0058), (**d**) TBK1_G217R_ (1.27±0.00, p=0.0002), (**e**) TBK1_R308Q_ (1.27±0.00, p=0.0182), (**f**) TBK1_R440X_ (1.33±0.02, p=0.0195), (**g**) TBK1_R440Q_ (1.18±0.01, p=0.0100), (**h**) TBK1_V464A_, (**i**) TBK1_M559R_, (**j**) TBK1_Q565P_, (**k**) TBK1_Q581H_ (1.10±0.00, p=0.005), (**l**) TBK1_Δ690-729_ (1.29±0.01, p=0.0034), (**m**) TBK1_E696K_ (1.38±0.04, p=0.0380) or (**n**) TBK1_I710N_ (1.16±0.02, p=0.0410). Images displayed are merged DAPI-stained nuclei (blue) and TDP-43 (yellow) channels. Nuclei and cell membrane outlines are defined by DAPI and TBK1 (FLAG) staining, respectively. Scale bars = 5 μm. (**o**) Quantification of the cytoplasmic/nuclear ratio of endogenous TDP-43 shows an increase in cells expressing some TBK1 variants (n=3 biological replicates of 250-300 cells each). Analysis done using repeated measures one-way ANOVA and Dunnett’s multiple comparisons test. Data is represented as mean ± SEM. *p<0.05; **p,0.01; ***p<0.001.

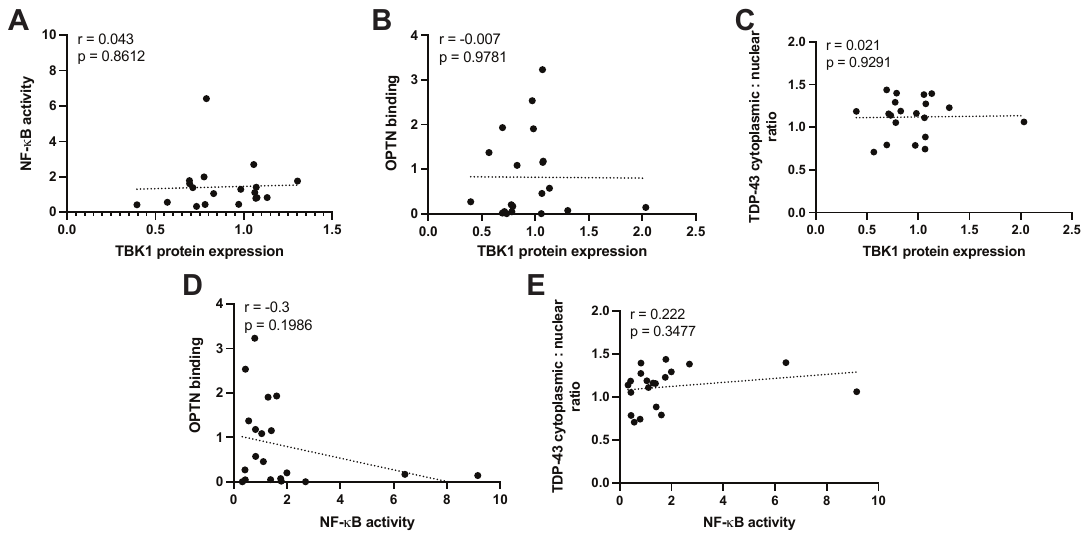

**Supplementary Figure 7.** Non-significant correlations between TBK1 functional assays. (**a**) Correlation analysis between TBK1 protein expression and NF-κB activity of TBK1 variants excluding one outlying variant. (**b**) Correlation analysis between TBK1 protein expression and degree of OPTN binding. (**c**) Correlation analysis between TBK1 protein expression and TDP-43 cytoplasmic mislocalisation. (**d**) Correlation analysis between NF-κB activity and degree of OPTN binding. (**e**) Correlation analysis between NF-κB activity and the TDP-43 cytoplasmic to nuclear ratio. Data is represented as mean ± SEM. r = Pearson correlation coefficient.

**Supplementary Table 1.** *TBK1* variants selected for functional studies.

| **Nucleotide** | **Protein** | **Disease** | **Minor Allele Frequency**  **(gnomAD v2.1.1, non-neurological cohort)** | **PolyPhen2** | **SIFT** | **Citations** | **Notes** |
| --- | --- | --- | --- | --- | --- | --- | --- |
| c.10A>G | p.T4A | FTD | 0 | probably damaging | damaging | [17] |  |
| c.77G>A | p.G26E | ALS | 0 | probably damaging | damaging | [17] |  |
| c.140G>A | p.R47H | ALS | 0 | probably damaging | damaging | [4] |  |
| c.281T>C | p.L94S | ALS | 0 | probably damaging | damaging | [4] |  |
| c.314A>G | p.Y105C | ALS, ALS-FTD | 0 | probably damaging | damaging | [4] |  |
| c.401G>A | p.R134H | ALS | 9.7 x10^-6^* | probably damaging | damaging | [7] |  |
| c.649G>A | p.G217R | ALS | 0 | probably damaging | damaging | [7] |  |
| c.829C>G | p.L277V | ALS | 1.15x10^-5^ | possibly damaging | tolerated | [7] |  |
| c.914T>C | p.I305T | ALS | 2.25x10^-5^ | probably damaging | tolerated | [4] |  |
| c.923G>A | p.R308Q | ALS | 0 | probably damaging | damaging | [4] |  |
| c.1135A>G | p.I379V | ALS | 8.73x10^-5^** | possibly damaging | damaging | [41] |  |
| c.1318-end deletion | p.△440-729 | ALS, FTD, ALS-FTD | - | - | - | [4,7,17] | mimics pathogenic nonsense mutation R440X |
| c.1319G>A | p.R440Q | ALS | 2.40x10^-5^ | probably damaging | damaging | [7] |  |
| c.1391T>C | p.V464A | - | 1.93x10^-2^ | probably damaging | tolerated | - | negative control: benign common SNP |
| c.1676T>G | p.M559R | ALS | 0 | benign | damaging | [4] |  |
| c.1694A>C | p.Q565P | ALS | 0 | probably damaging | damaging | [7] |  |
| c.1743A>T | p.Q581H | ALS | 0 | probably damaging | damaging | [7] |  |
| c. 2068-end deletion | p.△690-729 | - | - | - | - | - | artificial deletion of OPTN binding region |
| c.2086G>A | p.E696K | ALS, FTD, ALS-FTD | 1.15x10^-5^ | possibly damaging | tolerated | [4] | known pathogenic mutation |
| c. 2129T>A | p.I710N | ALS | 0*** | possibly damaging | damaging | [7] |  |

*5.60x10^-5^ in European (Finnish) population

**1.29x10^-4^ in Latino/Admixed American population

***2.17x10^-4^ in Other population
